# A nearest neighbour approach by genic distance to the assignment of individuals to geographic origin

**DOI:** 10.1101/087833

**Authors:** Bernd Degen, Céline Blanc-Jolivet, Katrin Stierand, Elizabeth Gillet

**Affiliations:** Thünen Institute of Forest Genetics, Sieker Landstrasse 2, 22927 Grosshansdorf, Germany; Forest Genetics and Forest Tree Breeding, Faculty of Forest Sciences and Forest Ecology, Georg-August-University of Göttingen, Büsgenweg 2, 37077 Göttingen, Germany

**Keywords:** Forensics, genetic assignment, genic distance, GeoAssign, geographic origin, timber, trees

## Abstract

During the past decade, the use of DNA for forensic applications has been extensively implemented for plant and animal species, as well as in humans. Tracing back the geographical origin of an individual usually requires genetic assignment analysis. These approaches are based on reference samples that are grouped into populations or other aggregates and intend to identify the most likely group of origin. Often this grouping does not have a biological but rather a historical or political justification, such as “country of origin”.

In this paper, we present a new nearest neighbour approach to individual assignment or classification within a given but potentially imperfect grouping of reference samples. This method, which is based on the genic distance between individuals, functions better in many cases than commonly used methods. We demonstrate the operation of our assignment method using two data sets. One set is simulated for a large number of trees distributed in a 120 km by 120 km landscape with individual genotypes at 150 SNPs, and the other set comprises experimental data of 1221 individuals of the African tropical tree species *Entandrophragma cylindricum* (Sapelli) genotyped at 61 SNPs. Judging by the level of correct self-assignment, our approach outperformed the commonly used frequency and Bayesian approaches by 15% for the simulated data set and by 5 to 7% for the Sapelli data set.

Our new approach is less sensitive to overlapping sources of genetic differentiation, such as genic differences among closely-related species, phylogeographic lineages and isolation by distance, and thus operates better even for suboptimal grouping of individuals.

## Introduction

In recent years, the application of forensic methods based on genetic markers to assign individual plants and animals to their geographic origin (Johnson *et al*, 2014; Ogden and Linacre, 2015) has gained importance for the control of trade regulations and consumer protection. Whereas many animal and plant species are protected by the Convention on International Trade in Endangered Species of Wild Fauna and Flora (CITES), the protection level can depend on the country of origin (species listed in CITES annex 3). For the timber trade, regulations such as the US Lacey Act and the EU timber regulation require the trader to declare the geographic origin of the material. Therefore it is important to be able to determine the true origin of the individuals that are traded (Johnson *et al*, 2014).

In different areas of the world, genetic reference data bases have been developed to aid in the identification of the true geographic origin of timber (Degen *et al*, 2013; Ng *et al*, 2016). To this end, reference samples of individual trees are collected throughout the natural distribution area of the species or in specific target regions. For each individual, the data base records the geographic coordinates of the individual and its genotype at gene loci representing different types of gene marker, especially molecular markers of the types nSSR, cp-DNA, mt-DNA and more recently SNP (Deguilloux *et al*, 2003; Ishida *et al*, 2013; Jolivet and Degen, 2012; Puckett and Eggert, 2016). Usually, more than one individual is sampled at each reference location, and reference locations and their sampled individuals are aggregated *a priori* into groups (or classes) by non-genetic criteria such as country of origin. The objective is to assign an individual of unknown origin, such as imported timber, to that group of reference individuals to which its multilocus genotype conforms best by some criterion. Examples for the assignment of individuals to place of origin have been published for timber tree species, elephants, and salmon (Degen *et al*, 2006; Glover *et al*, 2008; Wasser *et al*, 2015; Wasser *et al*, 2007).

Most of the previously applied approaches to the assignment of an individual of unknown origin to its putative group are frequency-based (Paetkau *et al*, 1995) and/or Bayesian (Rannala and Mountain, 1997). These approaches estimate the allele frequencies at all loci in every group, compute the probability that the multilocus genotype of the individual in question originates from each group, and assign the individual to that group for which the probability of origin is highest (Ogden and Linacre, 2015). Application of these approaches is based on assumptions that (a) the structure of the groups fits to Wright’s Island Model of drift (Wright, 1931), (b) the alleles at each locus are in Hardy-Weinberg-Proportions (HWP) (i.e., no stochastic association between alleles at the same locus), and (c) that the loci are in linkage equilibrium or are completely unlinked (i.e., no stochastic association between the genotypes between loci). Unfortunately, these assumptions are often violated in real populations. For example, molecular markers such as nSSRs often show deviation from HWP in at all spatial scales (Mariette *et al*, 2001; Nurtjahjaningsih *et al*, 2005). Also the use of large SNP marker sets derived from Next Generation Sequencing often leads to linkage disequilibrium (LD) in the data, which would require pruning process of loci in the data (Duforet-Frebourg *et al*, 2015). To overcome this problem and to avoid loss in the Gain of Informativeness for Assignment (GIA), other methods combine some loci into haplotypes (Duforet-Frebourg *et al*, 2015). Indeed, the use of haplotypes provides information on the ancestry and recombination events, which are useful when differentiation among groups is low. However, these methods are mostly interesting when dense marker sets are used (Gattepaille and Jakobsson, 2012), and it is not clear how these methods handle missing data, which is unfortunately the main issue with genotyping of timber or other material with degraded DNA.

The success of these assignment methods depends on the extent of genetic differentiation among the groups (Ogden and Linacre, 2015). The sampling scheme and the variability of the spatial genetic structure have an additional impact on the success of the assignment methods, especially if the groups represent political units such as countries.

When pre-defined aggregation to groups does not reflect the genetic structure among the reference individuals, genetic assignment approaches based on allele frequencies may fail (Meirmans, 2012). Grouping according to political (e.g. country) borders instead of genetic boundaries between populations as reproductive units is particularly critical and may lead to confusing genetic mixtures within groups (Manel *et al*, 2007). In the case that reference groups contain individuals of more than one reproductively isolated deme, such as different regions or even cryptic sympatric species, the mean genetic difference among individuals of different demes should be larger than the mean genetic difference among individuals of the same deme (Konnert *et al*, 2011). The ability of markers to identify such demes differs, however. For instance, the incomplete lineage sorting or chloroplast capture that can be observed among closely-related species at chloroplast markers (Duminil *et al*, 2013) could hinder attempts to recognize these species in the reference data. Also, the genetic differentiation among the groups of reference samples is biased by the proportion of species mixture or the mixing of individuals from different phylogeographic lineages (e.g. refugia) within the groups or by the assignment of individuals from the same spatial genetic unit (e.g. cross-border populations) to different groups.

The opposite approach to the *a priori* specification of groups is to attempt to partition individuals into reference groups that show HWP and linkage equilibrium. Bayesian clustering approaches, as implemented for example in the program STRUCTURE (Pritchard *et al*, 2000), have become common in recent years. These aggregations are, however, also based on estimates of allele frequencies that could be biased by unequal sample sizes among genetic groups or violation of the assumption that the populations fit Wright’s Island Model with relatively small, clearly differentiated populations (Meirmans, 2012). Another major drawback of the partitioning of reference individuals into genetic groups by Bayesian clustering methods is that the genetic groups they detect may be spread over more than one country. This collides with existing legislation that requires declaration of the country of origin, the political borders of which may cut through the middle of a genetic group, making it difficult to issue a statement from genetic testing on how likely this declaration is.

This paper describes a nearest neighbour classification approach that assigns unclassified individuals to predefined classes of reference individuals, such as by the country of origin that is of relevance for the timber trade or any CITES listed species with different country restrictions. The distance between individuals is measured by the genic distance between their multilocus genotypes, defining the nearest neighbours of a specific individual as those individuals with the smallest genic distance to it. An unclassified individual is assigned to a particular class if this class has the highest representation among a limited set of nearest neighbours by a new index *I*_*r*_ and if this representation is statistically significant. Since assignment is not based on estimation of allele frequencies within entire reference classes, the approach avoids the problems of possible discrepancy between political and genetic boundaries described above. We demonstrate its application using two data sets, a large set of simulated data for a hypothetical tropical tree species and experimental data of an African tree species. When applied to test whether individuals of known origin are correctly assigned to their class, it turns out that the probability of correct self-assignment is better than for conventional methods based on allele frequencies.

## Material and methods

### Nearest neighbour classification of individuals to origin

Consider the situation in which *R* reference individuals (index *j*) have been subdivided *a priori* into *G* different groups that correspond to their region (e.g. country) of origin. The objective is to assign each of a set of *T* test individuals of unknown origin to that group to which it is genetically most similar by the following procedure.

For a specified set of *M* gene loci, genetic similarity is first calculated on a pairwise basis between the test individual *i* and each reference individual *j* as the proportion of genes they share over the loci, that is, as 1−*D*_*ij*_, where

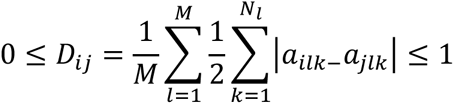

is the proportion of non-shared alleles (Gregorius, 1978). The index *k* denotes the alleles 1 to *N*_*l*_ at locus *l*. *a*_*ilk*_ is the frequency (or dosage) of the *k*-th allele at locus *l* in the test individual *i*, and *a*_*jlk*_ is the frequency of the same allele in reference individual *j*. For two diploid individuals, the frequencies of alleles at any locus can equal 1 (homozygote for that allele), 0.5 (heterozygote with one copy of the allele), or 0 (the allele is absent). *D*_*ij*_ is a direct measure of the genetic difference between two individuals. Compared to most other genetic distances, *D*_*ij*_ has the additional advantages that (a) it is a metric distance, (b) it ranges between 0 for the sharing of all alleles and 1 for the sharing of no alleles, and (c) it has a linear geometric conception (Gregorius, 1984).

All reference individuals *j* are then listed in ascending order of their genic distance *D*_*ij*_ to test individual *i*. If the number of loci is small, many reference individuals may have the same distance from the test individual, but as the number of loci is increased, the number of reference individuals with the same distance generally decreases until all distances are unique. Each number in the interval [0,1] specifies a neighbourhood of the test individual *i* as the set of all reference individuals *j* that have genic distance *D*_*ij*_ less than or equal to this number. Members of one neighbourhood are also members of any larger neighbourhood. The *k* nearest neighbours to test individual *i* by this measure are the *k* reference individuals that have the *k* smallest genic differences to test individual *i*.

In its most basic form, the *k*-nearest neighbour algorithm for classification would assign *i* to the group that has the highest number of reference individuals among its *k* nearest neighbors (“majority vote”). Its application here requires solution of the following problems: how to determine an “optimal” *k*, how to deal with groups containing different numbers of reference individuals, and how to test for significance of an assignment.

For consistency, the different numbers of reference individuals that have been sampled for different species, say, could be accommodated by specifying *k* as a recommended percentage *P* of all reference individuals. Alternatively, *k* could be specified as the number of reference individuals found within a neighbourhood of recommended size (critical genic distance), and test individuals whose critical neighbourhood is empty (*k*=0) would not be assigned to any group (outliers). Such a critical distance could also be chosen as the first “jump” in the cumulative frequency distribution of *1−D*_*ij*_ over the reference individuals *j*, separating genically similar reference individuals from genically dissimilar ones. For now, we will speak of the chosen individuals of highest genic similarity to the test individual as the “*k* nearest neighbours”.

Simple assignment of the test individual to the largest group among the nearest neighbours (“majority vote”) ignores the possibility that the original groups of reference individuals can be of different sizes, giving groups with more reference individuals a bigger chance to “win”. In order to compensate for group size, we define an index *I*_*r*_ that weights the relative frequency *o*_*r*_ (i.e., proportion) of each group *r* among the *k* nearest neighbours by the relative frequency *e*_*r*_ of group *r* among all reference individuals

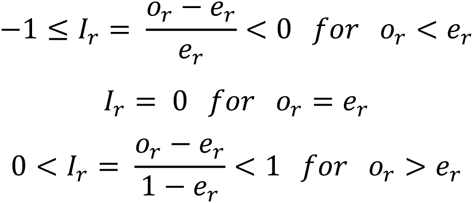

It holds that *I*_*r*_*=−1* whenever group *r* is not represented among the nearest neighbours, *I*_*r*_*= 0* if the frequencies *o*_*r*_ and *e*_*r*_ are equal, and *I*_*r*_*=1* whenever all nearest neighbours belong to group *r*. Unless the relative frequencies of all groups among the nearest neighbours are exactly the same as among all reference individuals, in which case *I*_*r*_*=0* for all groups (and *N* is divisible by *k*), at least one group *r* must have an index *I*_*r*_ greater than 0 and another group an index less than 0.

Selecting the group *r* with the highest value of *I*_*r*_ to be the major candidate for the origin of the test individual corresponds to the majority rule weighted by group size. The statistical significance of the higher relative frequency of group *r* among nearest neighbours compared to the set of all reference individuals can be tested by classifying reference individuals into two classes (i.e., members of group *r* and members of any other group) and calculating the P-value as the probability of finding at least the observed number *k*_*r*_*=k · o*_*r*_ of individuals from group *r* within a random sample of size *k* drawn without replacement from the set of all *N* reference individuals containing *N*_*r*_ individuals of group *r*, that is,

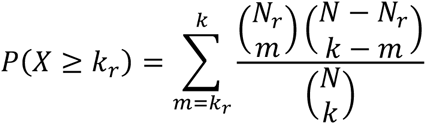

by the hypergeometric distribution, where 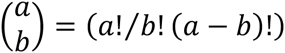 is the binomial coefficient. A P-value *P*(*X* ≥ *k*_*r*_) less than a chosen level of significance (commonly 0.05) is an indication that the observed representation of group *r* among the nearest neighbors is significantly high. It cannot be ruled out that another group with a smaller positive value of the index also shows significantly high representation among the nearest neighbours, making it an additional candidate for the origin of the test individual. This could happen, for example, if the test individual originates from a cross-border population. It must, however, be considered that hypergeometric tests of different candidate groups are not independent of each other.

By the same reasoning, a P-value *P*(*X*≥*k*_*r*_) greater than 0.95 indicates that this group has a significantly low representation among the nearest neighbours of the test individual and should be excluded as a candidate for the origin.

### Test of performance

We then compared the performance of this new individual genic distance assignment method with the so far commonly used genetic assignment methods based on frequency (Paetkau *et al*, 2004) and the Bayesian (Rannala and Mountain, 1997) approach criteria, referred below as frequency and Bayesian approaches.

As an indicator of the performance, we computed the proportion of correctly assigned individuals in self-assignments tests (Cornuet *et al*, 1999). Here the individuals of the reference data were self-classified to the sampled groups using the leave-one-out approach (Efron, 1983). For the frequency and the Bayesian approaches, we applied the self-assignment routine implemented in the program GeneClass 2.0 (Piry *et al*, 2004).

### Genetic diversity, Genetic differentiation and Hardy-Weinberg-Proportion

The success of genetic assignment depends on the level of genetic diversity, genetic differentiation among the groups and the similarity of the genotype frequencies to Hardy-Weinberg-Proportions.

As a measure for genetic diversity (effective number of alleles) we computed the mean allelic diversity v of group *r* (Gregorius, 1987):

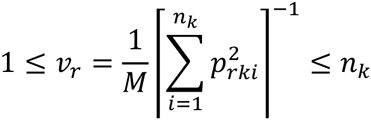

With relative frequency *p*_*rki*_ of the *i*-th allele at the *k*-th locus in group *r* and *n*_*k*_ different alleles at locus *k*.

To measure the difference between groups, we calculate the gene pool distance *d*_*0*_ between two groups *G*_*r*_ and *G*_*r′*_ as

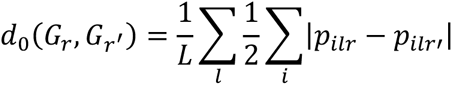

where the index *l* runs through the *L* loci, the index *i* runs through the alleles at locus *l*, and *p*_*ilr*_ and *p*_*ilr′*_ are the relative frequency of the *i*-th allele at locus *l* in group *G*_*r*_ and *G*_*r′*_, respectively (Gregorius, 1984). *d*_*0*_ ranges between 0 and 1. *d*_*0*_*=*0 holds when the relative frequency of every allele at every locus is the same in both groups. *d*_*0*_*=1* holds when the groups have no allele in common at any locus. To visualize difference between groups, cluster analysis based on the pairwise gene pool distances *d*_*0*_ between groups was performed using the Unweighted Pair Group Method with Arithmetic Mean (UPGMA) as implemented in the software PAST (Hammer *et al*, 2001).

Based on the gene pool distance, the complementary compositional differentiation *δ*_*SD*_ among the gene pools of the groups is then defined as the mean gene pool distance between each group *G*_*r*_ and its complement 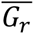, where 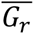 consists of all reference individuals that do not belong to group *r* (Gregorius, 1987), that is,

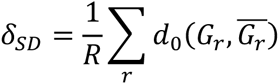

where the index *r* runs through the *R* groups, and 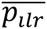 is the relative frequency of the *i*-th allele in the complement 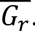. *δ*_*SD*_ ranges between 0 and 1. *δ*_*SD*_=0 holds when the relative frequency of every allele at every locus is the same in all groups. *δ*_*SD*_ = 1 holds when the groups have no allele in common at any locus.

For comparison, we also calculated the commonly used Wright’s FST (Wright, 1978) which is a measure of fixation (monomorphism) and not of the difference among groups (Gregorius *et al*, 2007).

As an indicator for the departure from Hardy-Weinberg proportions we computed the inbreeding coefficient *F*_*is*_ within each group *r* for which *He*_*rk*_ > 0 holds (Wright, 1950) as

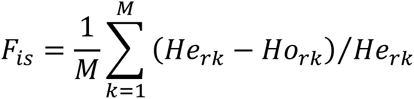

Here 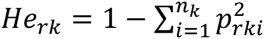 is the expected heterozygosity in group *r* at locus *k* under the hypothesis of random mating (i.e., the Simpson diversity) and *Ho*_*rk*_ is the observed heterozygosity (i.e., the relative frequency of heterozygotes).

### Simulated data

The simulation model “Eco-Gene Landscape” was used to create a baseline population for a tropical tree species in an area of 120 km by 120 km and its spatial genetic structure at 150 nuclear Single Nucleotide Polymorphism = nSNPs after 700 years of re-colonisation. The simulation model includes pollen and seed dispersal, mating, growth and mortality (Degen *et al*, 2006). The 120 km x 120 km landscape was subdivided into grid cells of side lengths 1 km 260 x 1 km (100 ha). Each grid cell represented a different population that is connected via pollen and seed dispersal with the surrounding populations. It was assumed that 60 % of the grid cells could be potentially occupied by the target tree species. This equals to 8640 populations in the simulated area. Simulation was initiated with 200 trees in each of only 10 grid cells to represent the refugia before re-colonisation. A mean individual growth rate of 0.5 cm diameter increment per year with a standard deviation of 0.2 was assumed. All trees with a diameter of 35 cm were fertile. Pollen dispersal was simulated following two normal distributions with means of 0 m and standard deviations of 500 m and 1500 m with a probability of 0.7 and 0.3, respectively. The resulting pollen dispersal pattern is typical for an insect pollinated species (Degen and Roubik, 2004). Random fertilization was assumed, with the exception that seeds from self-fertilization were inviable. Seed dispersal was also simulated by two normal distributions with means of 0 m and standard deviations of 100 m and 1500 m with probabilities of 0.7 and 0.3, respectively. Using these parameters, the majority of seeds were distributed within 300 m around the seed trees. Further, in accordance with field inventories typical for tropical trees, a maximum density of trees per ha in different diameter classes was considered. These maximum densities represented carrying capacities and caused mortality if by seed production, migration or ingrowth the densities were exceeded. The model thus created overlapping generations.

A total of 888,841 individuals were generated over all populations during 700 simulated years. At that time, 6212 grid cells contained populations. The number of individuals in each population varied from 2 to 206 with an average of 156. We assumed that every SNP locus has 2 alleles (A1, A2). The initial allele frequencies were randomly selected with 0<A1<1 for all initial populations (i.e., uniform distribution). To create reference data, we selected 10 areas, each with a size of 10 km by 10 km (Figure 1). In each of these areas we randomly selected a reference data set of 100 individuals. In order to imitate the effect of species mixture, in area 5 we replaced allele “2” by an allele “3” in all individuals, and in areas 6 and 7 we did the same for 50% of the individuals. Thus the areas 1, 2, 3, 4, 8, 9, and 10 are genetically purely species A, area 5 is purely species B, and the areas 6 and 7 are mixtures of 50% species A and 50% species B.

**Figure 1:**
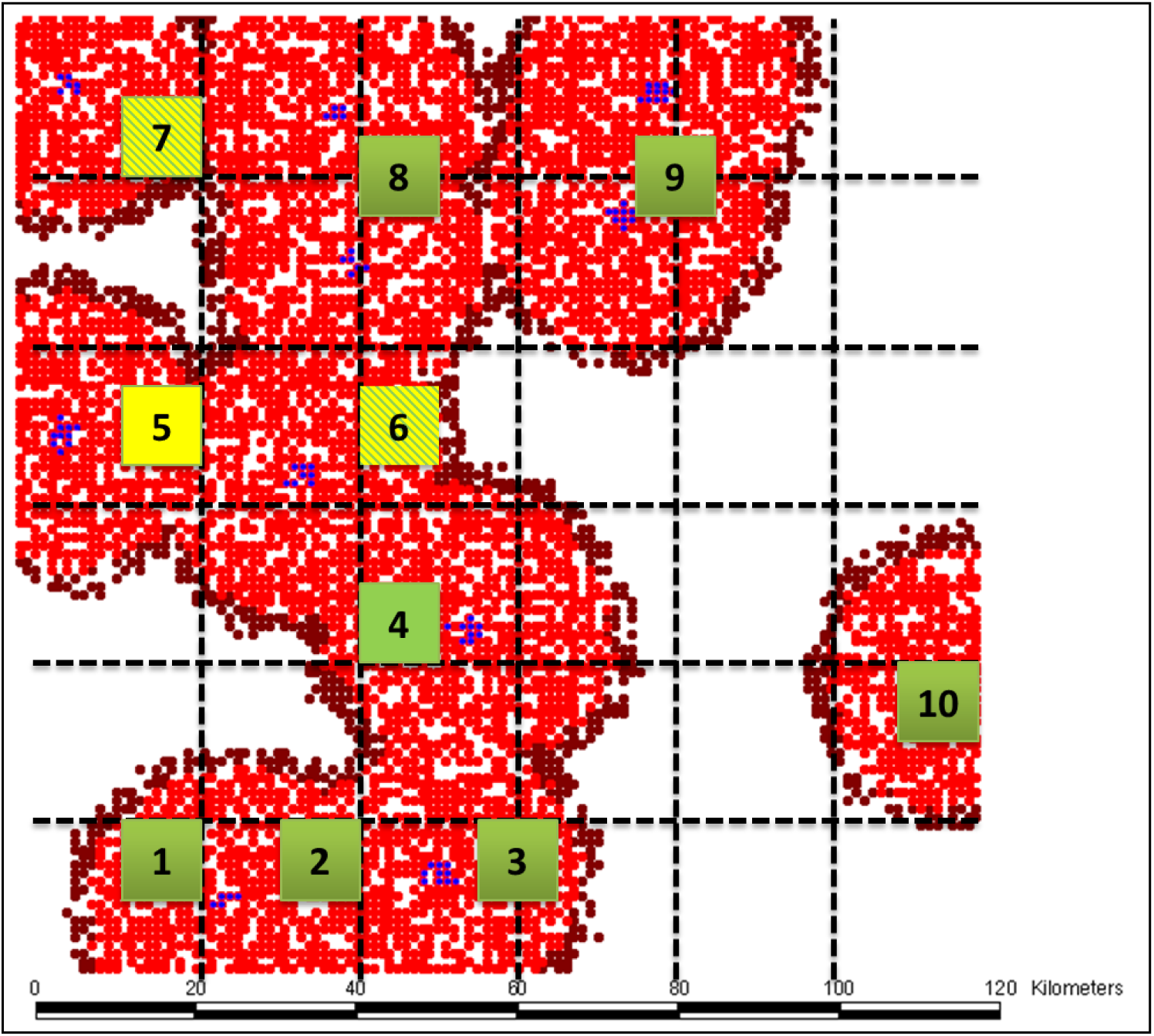
Simulated data set; red and brown points are grid cells covered by trees after 700 simulated years of re-colonisation. Blue points = location of initial refugial areas. Green squares mark selected areas of species A, yellow squares mark selected areas of species B, and yellow-green squares are selected areas with 50% mixtures of species A and B

### Experimental data on Sapelli

Sapelli is the trade name of the botanical species *Entandrophragma cylindicum* of the same family as mahogany (Meliaceae). Since decades, this high value timber tree is one of the most harvested tree species in the natural forests of tropical Africa. In frame of a project of the International Tropical Timber Organization (ITTO), we developed a genetic reference data set in order to check claims on the geographic origin of the traded timber (Degen and Bouda, 2015). A total of 1292 Sapelli trees were collected in seven African countries (Figure 2). Then, SNPs were developed and all individuals were screened at 61 SNPs. Two Sapelli samples from Cameroon and DRC were used as reference for RAD (Restriction Site Associated DNA) sequencing and SNP detection (Pujolar *et al*, 2014). A total of 140 putative SNP loci were screened with a MassARRAY technology (Agena Biosciences, San Diego, California, USA) on a set of 190 reference individuals chosen from different countries among the reference samples (McKernan *et al*, 2002). Estimates of genetic differentiation and correlations among genetic and geographic distances per locus were used together with amplification success rates to select 61 loci for reference sample screening. Details are presented elsewhere (Blanc-Jolivet and Degen, in preparation).

**Figure 2:**
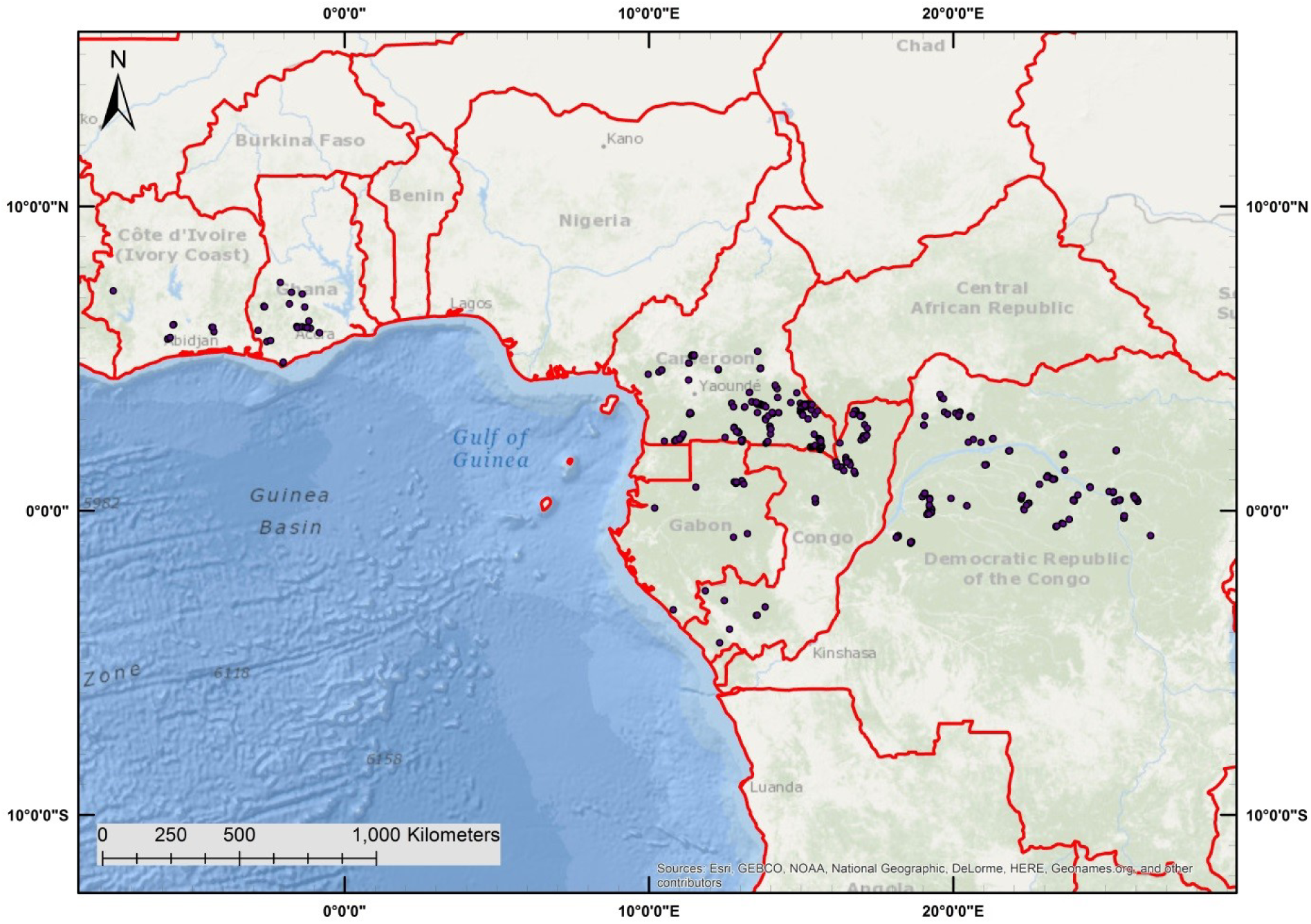
Location of the sampled trees of *Entandrophragma sp*. in Africa

## Results

### Simulated data

After 700 years of simulated re-colonisation, the allelic diversity (*v*_r_) in the selected areas varied between 1.52 and 1.58 for the individuals belonging to species A and in the same range for individuals of species B. The fixation index equalled F_ST_ = 0.053 and the compositional differentiation *δ*_*SD*_=0.082 for species A and FST = 0.045 and *δ*_*SD*_=0.106 for species B. For all individuals of both species together, the measures among the 10 selected areas equalled FST = 0.171 and *δ*_*SD*_=0.196. Both measures clearly indicated a stronger fixation and differentiation between the species compared to the within species differentiation.

Spatial-genetic analysis reveals a clear spatial structure within species A showing that individuals up to a distance of 20 km are genetically more similar than expected for a random distribution (Figure 3).

**Figure 3:**
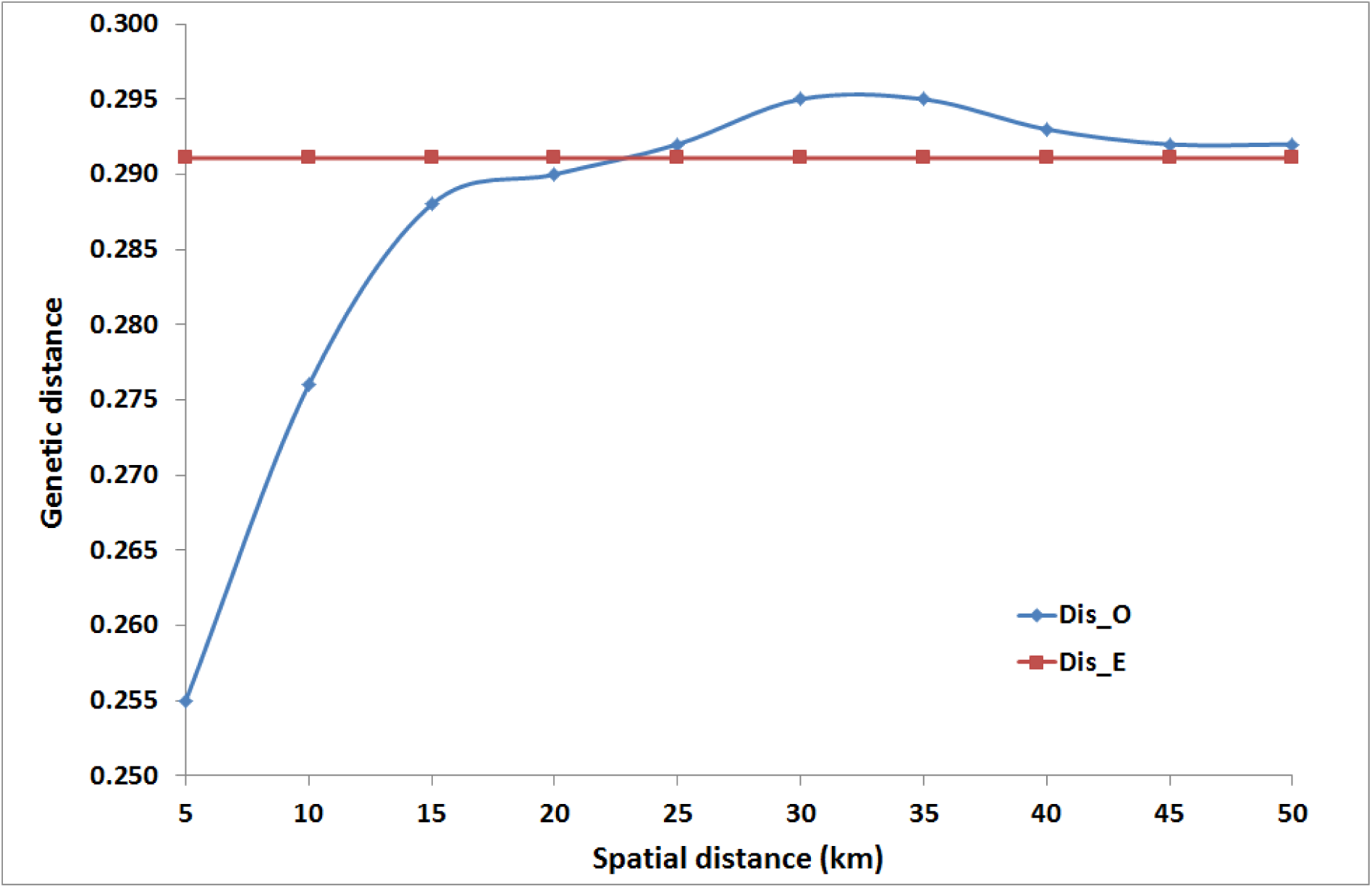
Spatial genetic structure of the simulated data set for species A. Dis_O is the observed mean genic distance among individuals in the given spatial distance class. Dis_E is the expected genic distance for a random distribution of individuals in space.

The dominating species signal is visible in the genetic differentiation among the areas (Figure 4). The areas covered only by species A (green) form one group in the dendrogram and the areas with pure species B (yellow) and species mixture (light green) form another group.

**Figure 4:**
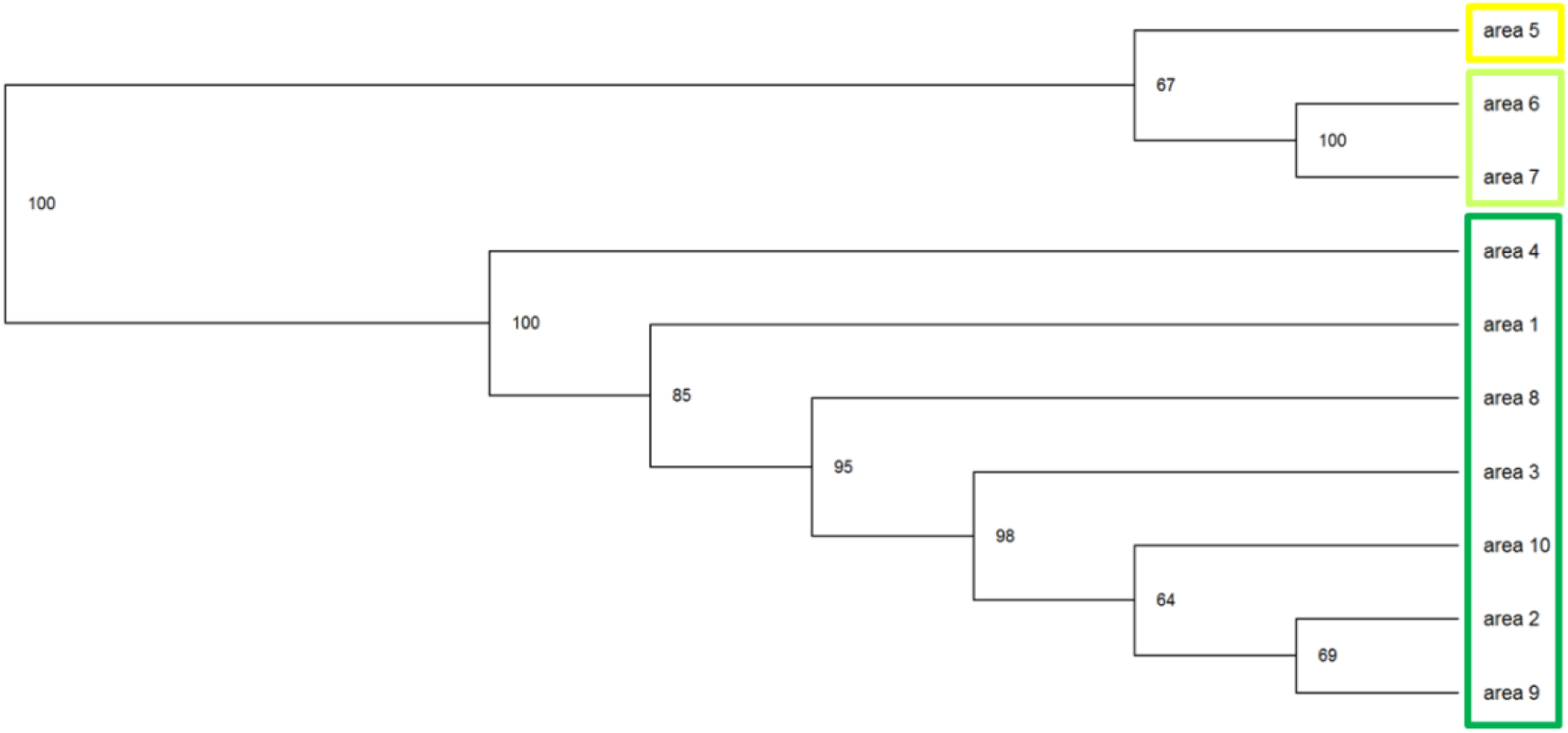
Dendrogram based on a cluster analysis (UPGMA: Unweighted Pair Group Method with Arithmetic Mean) using the genic distance matrix among the allele frequencies of the ten selected areas in the simulated data set. The numbers are the proportions of the 1000 bootstrap simulations for shifting the included SNP loci that find the same branch.

In addition, results for the self-assignment is quite stable over a range of percentiles of nearest neighbours (P= 0.5% to 10%, Figure 5).

**Figure 5:**
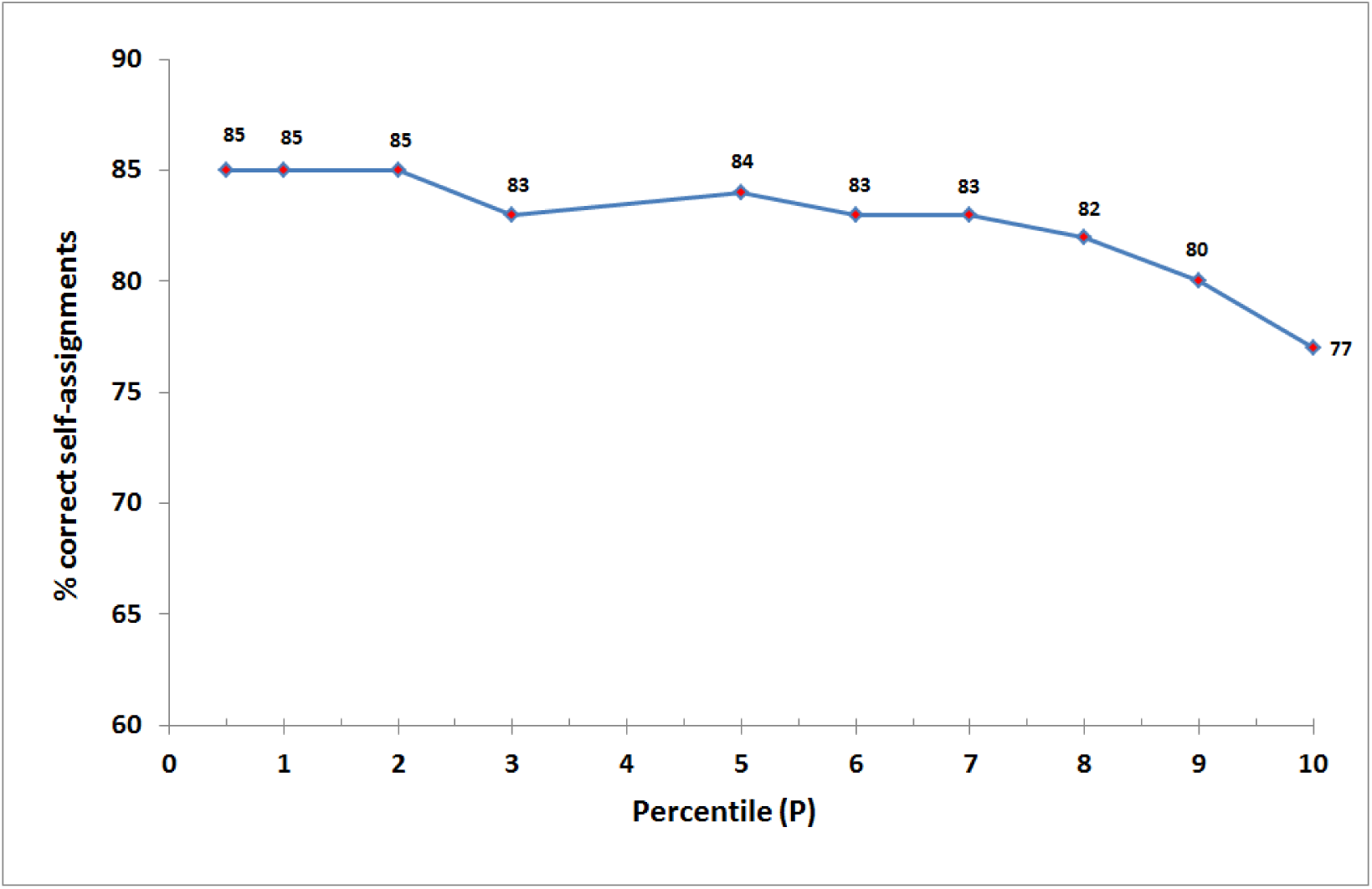
Proportion of correct self-assignments for different percentiles (P) of the genically most similar reference individuals (nearest neighbours) in the simulated data set

The frequency and the Bayesian approach assigned 70% of all 1000 individuals to the correct area, whereas our nearest neighbour approach assigned 85% of the individuals correctly (Table 1). The largest differences were observed for the areas 6 and 7 with the simulated species mixture of individuals. There, the frequency and the Bayesian approach did not assign a single individual correctly but the nearest neighbour approach accurately assigned 95% for area 6 and 82% for area 7. The frequency and Bayesian approach performed slightly better than the nearest neighbour approach in areas (areas 2, 3, 9) with lower genetic differentiation (lower values of *δ*_*SD*_) and heterozygosity values close to Hardy-Weinberg-Proportion (Fis values close to 0), whereas our nearest neighbour approach came up with better results for areas with stronger genetic differentiation and stronger departure from Hardy-Weinberg-Heterozygosity (areas 1, 6, 7, 8).

**Table 1:** Results of the self-assignments for the simulated data set using the different assignment approaches, N = sample size, *δ*_*SD*_ = complementary compositional differentiation, F_is_ = mean inbreeding coefficient over all loci

| Area | N | $\delta_{SD}$ | $F_{is}$ | Nearest neighbour approach (P1) % | Frequency approach % | Bayesian approach % |
| --- | --- | --- | --- | --- | --- | --- |
| Area 1 | 100 | 0.170 | 0.026 | 99 | 93 | 94 |
| Area 2 | 100 | 0.142 | 0.040 | 67 | 77 | 77 |
| Area 3 | 100 | 0.147 | 0.027 | 81 | 91 | 91 |
| Area 4 | 100 | 0.174 | 0.035 | 93 | 95 | 96 |
| Area 5 | 100 | 0.463 | 0.029 | 100 | 100 | 100 |
| Area 6 | 100 | 0.228 | 0.340 | 95 | 0 | 0 |
| Area 7 | 100 | 0.197 | 0.304 | 82 | 0 | 0 |
| Area 8 | 100 | 0.156 | 0.057 | 87 | 84 | 85 |
| Area 9 | 100 | 0.140 | 0.013 | 67 | 78 | 78 |
| Area 10 | 100 | 0.143 | 0.014 | 74 | 79 | 79 |
| Total / Mean | 1000 |  |  | 85 | 70 | 70 |

### Experimental data on Sapelli

The samples from Ghana and Ivory Coast were merged in one group because of the small sample sizes and the geographic neighbourhood. The mean allelic diversity (v_r_) varied between 1.39 and 1.59 in the different countries. The mean fixation was FST = 0.092 and the mean compositional differentiation *δ*_*SD*_=0.126 (0.078−0.194). There was a clear spatial structure showing that individuals up to a distance of 500 km are genically more similar than expected for a random distribution (data not shown). The analysis with STRUCTURE came up with a clear indication that a significant proportion of individuals represent other species within the genus *Entandrophragma* (Blanc-Jolivet and Degen, in preparation).

The frequency and the Bayesian approach assigned 68% and 66% of all individuals to the correct area, whereas the nearest neighbour approach assigned 74% of the individuals correctly (Table 2).

Large differences among the methods were observed for individuals from Cameroon. Here, the nearest neighbour approach assigned 77% of the individuals correctly, while the frequency and the Bayesian approach assigned only 65% and 61% correctly. This was the group with the largest sample size. Also for samples from DRC with the second largest sample size the nearest neighbour approach was better.

As for the simulated data, the nearest neighbour approach was better in groups with high genetic differentiation and stronger departure from Hardy-Weinberg heterozygosity (DRC, Ghana/ Ivory Coast) and the other approaches performed better in situations with lower differentiation and F-values close to 0 (Gabon).

**Table 2:** Results of the self-assignments for the Sapelli data set using the different assignment approaches, all data that had at least 50% of the loci were included. N = sample size, *δ*_*SD*_= complementary compositional differentiation, F_is_ = mean inbreeding coefficient over all loci

| Country | N | $\delta_{SD}$ | $F_{is}$ | Nearest neighbour approach (P=1) % | Frequency approach % | Bayesian approach % |
| --- | --- | --- | --- | --- | --- | --- |
| Cameroon | 430 | 0.094 | 0.234 | 77 | 65 | 61 |
| Congo Brazzaville | 140 | 0.096 | 0.269 | 44 | 51 | 50 |
| DRC | 419 | 0.170 | 0.439 | 86 | 81 | 78 |
| Gabon | 36 | 0.078 | 0.233 | 3 | 19 | 22 |
| Ghana / Ivory Coast | 89 | 0.194 | 0.312 | 74 | 71 | 67 |
| Total/ Mean | 1114 |  |  | 74 | 68 | 66 |

## Discussion

### Realistic simulated data and typical large scale SNP data set of a tree species?

So far, not many data sets have been published on large scale genetic differentiation of tree species at SNP gene markers, and most of them are looking for SNP loci linked to selection (Csillery *et al*, 2014). The large scale genetic structure at the 150 nuclear SNPs in the simulated data was the result of non-selective processes (gene flow, mating system) and the imitated effect of a mixture of two closely related species. The level of genetic differentiation and fixation we observed in the Sapelli data set (*δ*_*SD*_ = 0.123, F_ST_= 0.092) lay between the pure within-species differentiation and fixation (*δ*_*SD*_ = 0.083, F_ST_= 0.053) and the overall genetic differentiation and fixation of the simulated data set (*δ*_*SD*_ = 0.196, F_ST_=0.171). A similar level of fixation (F_ST_=0.15) has been reported for a large SNP data sets of *Populus trichocarpa* in North America (Slavov *et al*, 2012). The overlapping of different genetic signals such as species hybridisation and mixture, phylogeographic pattern and isolation-by-distance effects responsible for the genetic differentiation of tree populations has been found in several recent studies (De La Torre *et al*, 2015; Jelic *et al*, 2015; Thomson *et al*, 2015). We imitated the species signal by replacing one of two alleles by a third allele, keeping one allele at each locus in common among species. This might look rather “artificial”, but in nature separation of related species of the same genus is visible by the same observation (mixture of species-specific alleles and common alleles (see e. g. for different citrus species (Curk *et al*, 2015) and for different frog species (Lemmon and Lemmon, 2012)).

### Performance of assignment methods

We demonstrate by the simulated and the real data set of the African tree species Sapelli that our nearest neighbour approach had a higher percentage of correctly assigned individuals compared to the frequency and Bayesian methods. The reason for this better performance was that our calculated index takes into account only the genically most similar individuals to the reference data. Thus, it reduces the bias of overlapping genetic signals because there is a hierarchy of genic similarity (Mori *et al*, 2015; Petit *et al*, 2005). Individuals of the same species (or the same level of hybridisation), from the same phylogeographic lineage and the same geographic region are the most similar. On the opposite side, individuals from different species show the highest genic differences. Of course, one could use Bayesian clustering analysis to obtain a genetic grouping of reference data that minimizes departure from HWP within each group (Pritchard *et al*, 2000), thereby limiting problems due to population sub-structure. But for many applications – especially as part of law enforcement and forensic investigations – the investigator needs to come to a conclusion for a given (non-biological) grouping of reference individuals. For example, the question for application of the EU-timber regulation is to confirm or not that the timber is from a Sapelli tree grown in Cameroon. In this case, the data must be aggregated by countries. The better performance of the nearest neighbour approach was particularly visible for groups with high sample sizes, higher genetic differentiation and an excess of homozygotes compared to Hardy-Weinberg expectations. The excess of homozygotes could be caused by a subpopulation structure within the groups (Wahlund effects). The power of our nearest neighbour approach depends on a sufficient sample size from all of the different subpopulations represented in the reference data. On the other hand, there were also groups within these data sets for which the Bayesian and the frequency approach had higher success rates for the self-assignments. These were groups with lower genetic differentiation and heterozygosity close to Hardy-Weinberg-expectations.

In our individual distance approach, a decision is still needed on the percentile of genically most similar individuals to be included in the calculation of the index. Thus we need to decide on the right number of nearest neighbours. We have shown with the simulated data set that the percentage of correctly assigned individuals does not change much for percentiles from P=0.5% to P=5% (Figure 5). For all approaches, the success of the assignment depends on the genetic differentiation among the reference individuals (Ogden and Linacre, 2015). If there is only little genetic differentiation, then it is better to include more individuals in the calculation of the index because the power of the statistical test increases with increasing numbers of reference individuals included. On the other side, large percentiles will increase the mixture of genic distances caused by different processes.

Cornuet *et al* (1999) compared the performance of different assignment methods using simulated data sets. They found that the Bayesian and the frequency approaches outperformed by far different genetic distance approaches. Our study found the opposite. The difference is that Cornuet *et al* (1999) included all individuals of each population in the analysis, and their simulated data set fulfilled the assumptions of Hardy-Weinberg and linkage equilibrium. Our simulated data set and the Sapelli experimental SNP-data were characterised by an excess of homozygotes in nearly all groups. This excess could be explained by Wahlund effects, thus by a mixture of individuals from different populations in the same groups (Wahlund, 1928).

### Forensic application in timber tracking

We have generated and are continuously developing genetic reference data for timber tracking of several high-value tree species (Degen and Bouda, 2015; Degen *et al*, 2013). A few years ago, we shifted from SSR markers to SNPs because their shortness is an advantage for amplification in the typically degraded DNA of timber (Jardine *et al*, 2016; Pakull *et al*, 2016). These tools are developed to reduce illegal logging. As has been pointed out by Ogden and Linacre (2015), wildlife forensics is faced with several drawbacks compared to the ideal situations found in human forensics. It is often a big challenge to obtain a collection of reference material that is well distributed over the geographic range of the target tree species. It is difficult to obtain access to many remote areas and sometimes even dangerous because of unstable political situations in countries where illegal logging is an issue. We often need to collaborate with local groups that help with the collection of material. They receive training but are usually not scientists. Yet even for the botanical specialist it is often impossible to clearly distinguish between closely related tree species based on morphological traits. Thus it is not uncommon for us to have reference samples that contain admixtures of closely related species. This is either due to errors in the field work or the lack of an efficient biological barrier between closely related species. This underlines the importance of assignment routines that are robust against this kind of error and still yield reliable results.

## Access to the program

We implemented the approach in a software application called GeoAssign and this is available through a web service at https://geoassign.thuenen.de. The webserver architecture is designed around a Ruby on Rails application that handles multiple jobs via a Sidekiq (http://sidekiq.org) / Redis (http://redis.io) background processing framework. The GeoAssign method itself is implemented in the Ruby programming language and included in the web application as a gem.

## Outlook

We are currently extending our nearest neighbour approach using a city-block distance for metric traits such as stable isotopes. The idea is to have a single statistical assignment approach for different data sources that should be combined to increase the performance of forensic assignment approaches (Dormontt *et al*, 2015; Lowe *et al*, 2016). Applying the same assignment routine would also enable us to better compare the performance of different reference data sets (such as genetic, stable isotope or near infrared data).

As is the case for the other assignment approaches, the nearest neighbour approach will always assign the test individual to a group. But it is possible that the true group of origin is not represented by the reference individuals. To deal with this problem, other approaches simulate genotypes based on allele frequencies in the different groups of reference data, compute the distribution of likelihoods for assignment of these simulated genotypes to “their” group, and compare the likelihood of test individuals to these distributions and interpret the position in the distribution as an “exclusion probability” (Cornuet *et al*, 1999). Again this approach suffers from the assumptions to be made on the use of allele frequencies to simulate genotypes. To deal with these “alien individuals”, we are integrating a routine into our nearest neighbour approach that compares the genetic distance of the test individuals to the distribution of genetic distances among all reference individuals within each group. An outlier would be identified by its relatively large genic distance compared to the distribution of distances among members of the same group.

## Acknowledgement

The work of CB-J was financially supported by the International Tropical Timber Organization (ITTO) through the project PD 620/11 Rev.1 (M): “Development and implementation of species identification and timber tracking in Africa with DNA fingerprints and stable isotopes”. The work of EMG was financially supported by grant ZI 662/7-1 of the Deutsche Forschungsgemeinschaft. Further we thank Malte Mader for the setup and configuration of the LINUX web server. Finally, we are thankful to the two reviewers for their very useful comments and suggestions on a former version of the manuscript.

